# Evolution of multicellularity and unicellularity in yeast *S. cerevisiae* to study reversibility of evolutionary trajectories

**DOI:** 10.1101/2020.08.15.252361

**Authors:** Phaniendra Alugoju, Anjali Mahilkar, Supreet Saini

## Abstract

Adaptive trajectories of populations have been focus of number of studies. However, adaptive trajectories have not been studied in the context of reverse evolution. By reverse evolution, we mean a scenario where selection is reversed. In this work, we use evolution (and reversal from) of multicellularity in *S. cerevisiae* as a model to answer this question. When selected for fast-settling variants, multicellularity evolves rapidly in the organism. On reversing selection, unicellularity evolves from the multicellular clusters. However, the dynamic trajectories of the two processes are different. In this context, evolution is not reversed dynamically at a phenotypic level. The phenotypic reversal is not driven by reversal of the original mutations during the forward evolution. Overall, our results show that the dynamics of molecular and phenotypic trajectories of evolution are distinct, and reversal of selection leads to unique trajectories of phenotypic reversal.

## Introduction

Reversibility, in an evolutionary context, is often seen as return to ancestral state (1). Historically, investigation of reversibility of evolutionary processes has been an important theme in study of evolution (2–5). In microbial systems, this question has been addressed in a number of experimental studies, and has led to different answers (6, 7). In these experiments, a population is shifted from one environment to another, and adapted for a certain number of generations. The evolved population is then shifted back to the original environment, and its adaptation studied. In this context, reversibility is studied as the ability/dynamics of a population to re-adapt in an environment, when shifted from another (7). Towards this end, one of the most well studied systems has been evolution of sensitive bacteria in a population of resistant bacteria, when a stressor (antibiotic/phage) is removed from the environment (8–10). In the context of microbial systems, sequencing makes it possible to trace the reversibility of evolution at genetic or molecular level. In bacteria, when antibiotic resistance was removed, the original genotype was not restored (11, 12). On the other hand, evolution experiments with virus have demonstrated that phenotypic reversals can map to precise nucleotide reversals (6, 13). Enzymes, too, have been shown to recover activity, albeit via different mutational paths (14). The broad lesson from these experiments is that the longer the time spent in the novel environment, the more difficult the likelihood of reverse evolution becomes, when the population is re-introduced in the original environment.

However, reversibility in another context, has been largely unexplored. What happens to the evolutionary fate of a population, when the selection pressure is reversed? In this context, to study reversibility of evolution, we need systems where an exact and opposite selection pressure is applied on a system. Such a choice of a system is not trivial. For instance, is absence of an antibiotic the reverse of evolution in presence of (how much?) antibiotic? Or, is evolution in presence of glucose the reverse of evolution in glycerol? To answer this question, we study evolution (and reverse) of multicellularity in yeast, using settling (or lack of) as a selection force.

The evolutionary transition from single cells to multicellular organisms was one of the major events in the history of life on Earth (15). Eukaryotic multicellularity has evolved independently at least 25 times in distinct lineages (16–18). However, understanding the evolution of complex multicellular organisms from unicellular organisms is challenging, because the first steps in this process occurred in the deep past (16, 19). Mathematical models have demonstrated key steps involved in the transition from unicellular form to multicellular clusters (20–23). Experimentally, yeast *Saccharomyces cerevisiae* has been used for studying the origin of multicellularity in eukaryotes (24–32). These studies have demonstrated that under centrifugal selection, the transition from unicellularity to multicellularity is relatively quick. The mechanism of cluster formation in yeast is through post-division adhesion, but not through aggregation (24). This de novo origin of snowflake yeast clusters is mainly due to mutations in transcription factor ACE2 gene (33, 34). This method of growth ensures high relatedness among component cells of a multicellular cluster, therefore reducing internal conflicts. Clusters reproduce by fragmentation especially when tension among cells in the cluster exceeds the tensile strength of cellular adhesion (25). Apoptosis, a major determinant in the development and somatic maintenance in metazoans, is also involved the reproduction of yeast multiclusters (34).

Though largely unicellular, there are some yeast strains which exit as multicellular clumps in their ecological niches. The multicellular existence is thought have been driven because of the benefit it provides under conditions of stress (35). These clusters, when compared with the unicellular counterpart, were found to be more resistant to oxidative and solvent stresses. This increased stress resistance however came at a cost of lower growth rate, in the absence of stress.

Depending on the condition, yeast can show bidirectional transitions between unicellularity and multicellularity (30, 36–38). A recent study showed that unicellular yeast evolve from wild type clump-forming ancestors by propagating samples from suspension after larger clumps have settled (35). Thus, yeast clusters serve as an excellent model for studying the dynamics of reverse evolutionary events.

In this work, we first evolved yeast multicellular clusters from unicellular yeast *S. cerevisiae* using settling as a selection force. The multicellular cluster was verified to be because of mutation in ACE2 gene, and not simply due to cell-cell adhesion. We show that the multicellular clusters perform better than the ancestor in presence of various stresses. The multicellular cluster is then reverse evolved. Would reversal of selection lead the population to re-trace its path on the phenotypic space to the ancestral phenotype? Or, would availability of a large number of genetic solutions, in response to a precisely opposite environmental selection, dictate that the reverse path is largely unpredictable? Our results from the reverse evolution experiment show that, at the phenotypic level, the kinetics of evolution of unicellularity does not simply retrace the path that was taken to multicellularity. The genetic solutions that help accomplish this change are also distinct, and not because of mutations in ACE2. Study of bidirectional transitions between multi- and unicellularity thus offer a model system to study reversibility in an evolutionary process.

## Materials and Methods

### Yeast strain and growth medium composition

Yeast *Saccharomyces cerevisiae* haploid strain SC644 (*MATa MEL1ade1 ile trp1-HIII ura3-52*) was used (39). Cells were grown in test tubes (25 x 150 mm) containing liquid YPD (1% yeast extract, 2% peptone and 2% glucose) medium at 30°C. Unless stated otherwise, all cultures were grown in an orbital shaker at a speed of 250 rpm for 24 h. Yeast extract and Peptone were procured from Himedia, Mumbai, India. Glucose was procured from Sigma Aldrich, USA. All the chemicals used in this study were of highest analytical grade.

### Selection for multicellularity

Multicellular yeast clusters were generated as described by Ratcliff et. al. 2012 (24) (Figure 1A). A single isogenic yeast colony was inoculated into 10 ml of YPD medium and incubated at 30°C with 250 rpm for 24 h. The resulting culture was then used as a pre-inoculum for the initiation of the selection for multicellularity. An aliquot (1 ml) of overnight grown yeast culture was transferred to a 1.5 ml microcentrifuge tube and then centrifuged at a speed of 100g for 15 sec. The bottom 100 μL was transferred to a tube containing fresh YPD media and allowed to grow (30°C, 250 rpm) for 24 h. This process was continued for 60 transfers. Six independent lines under centrifugal selection pressure and the control were studied in this work. A control line was evolved in YPD, from which, 100 μL cultures was transferred into fresh YPD after growth for 24 h (30°C, 24 h). In this control, the transfer was immediate without allowing the cells to settle down. Glycerol freezer stocks, in 30% glycerol, were made for every tenth transfer and stored at −80°C.

**Figure 1.**
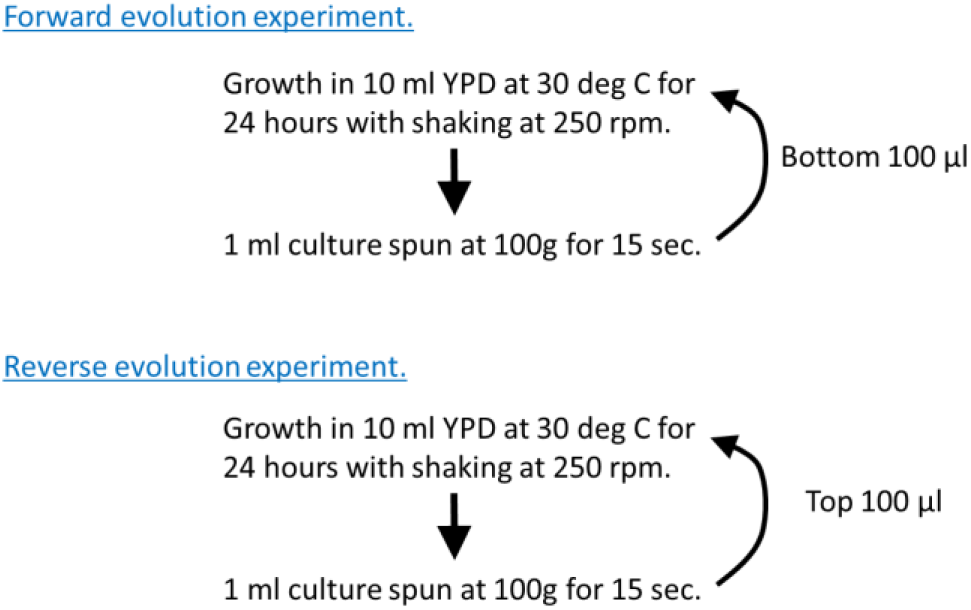
**(Top)** The experimental evolution of multicellularity under centrifugal selection pressure. **(Bottom)** Experimental plan for reverse evolution of unicellularity from multicellularity under centrifugal selection pressure.

### Selection for unicellularity

The frozen glycerol stocks of multicellular clusters obtained after 60 transfers were streaked onto YPD plates and incubated at 30°C for 48 h (Figure 1B). Cells from a single colony were spread on glass slide and a cover slip. Images were taken using a microscope (Labomed Binocular) under 40X magnification. A single multicellular cluster was then inoculated into fresh YPD medium and incubated at 30°C for 24 h with shaking. These cultures were used as ancestors for the reverse selection of unicellular phenotype from the multicellular cluster phenotype. As described above, an aliquot (1 ml) of overnight grown yeast culture was transferred to centrifuge tubes and centrifuged at a speed of 100xg for 15 sec. In this experiment, the top 100 μL was transferred into tubes containing fresh YDP medium every 24 h. Glycerol stocks were made for every tenth transfer in 30% glycerol and stored at −80°C.

### Characterization of colony phenotype

Each of the overnight grown (24 h) yeast population streaked onto YPD plates. These plates were then incubated at 30°C until single colonies appear, at which point, images of the plates were taken (28).

### Growth experiments

Saturated yeast cultures, from both forward and reverse evolution experiments, were diluted in fresh media to get an initial optical density at 600 nm wavelength (OD600) to 0.1. The diluted culture was allowed to grow at 30°C and growth kinetics monitored in a Tecan Infinite M200 Pro microplate reader. The microplates were overlaid with a Breathe Easy membrane (Sigma).

To test growth of yeast clusters in ethanol stress, overnight cultures were diluted in fresh media to an OD600 of 0.1. Ethanol (to a final concentration of 0, 2.5%, 5%, and 10%) was added to the cultures and the cell growth was checked by measuring the OD600 using a 96-well plate reader. OD600 values were taken every one hour over a 30 h period.

### Measurement of yeast cluster size

Yeast populations from both forward and reverse evolution experiments were grown in 10 ml of fresh YPD media shaken at 250 rpm for 24 h. These overnight cultures were transferred to fresh YPD medium and grown until exponential phase (OD600 ~ 2). About 100 μl of exponentially growing cells were washed thrice with PBS and analyzed for FSC-A of each cells/cluster using flow cytometer. A minimum of 30,000 clusters per population were analyzed and the average of the FSC-A at the indicated tome of transfer time was calculated (40).

### Settling assay

Yeast cultures from forward evolution experiments were inoculated from single colonies into 10 ml YPD liquid cultures and grown overnight at 30°C. Each culture tube was vigorously vortexed and placed on a lab bench at room temperature to allow for settling. Images of each population during the settling were taken at different time intervals (0, 15, 30, and 60 min) (41).

### Stress resistance assay

Each of the yeast populations was grown at 30°C with 250 rpm shaking for 20 h. Cells were serially diluted to OD600 of 3, 1, 0.3 and 0.1. A 4 μl volume of these cultures was spotted onto YPD plates. In order to test the effect of cold and heat stress, a set of potted plates were also incubated at 20°C and 37°C, respectively. A single plate was incubated at 30°C as a control. Images of the plates were taken at different days of incubation.

### Cloning of ACE2 gene in pT1-3B plasmid

ACE2 promoter region and the coding sequence was amplified from the genomic DNA of the ancestor using the primers (5’GGG GTC GAC TTA GTT AAC TCT ATC TAT TG3’) and (5’GGG AAG CTT TTT TTG GCC CTT AAG ACT AC3’). The PCR product and the plasmid pT1-3B (42) was digested with the restriction enzymes Sal1 and HindIII (New England Biolab (NEB)). T4 DNA ligase was used to construct the resulting plasmid pACE2, in which ACE2 expression was under its native promoter.

In addition to the ACE2 sequence from the ancestor, two other templates were used to construct plasmids pACE2T40 and pACE2T50. For the first (pACE2T40), the ACE2 promoter and coding region was amplified from the genomic DNA of cells in line 1 after 40 transfers. For the second (pACE2T50), the ACE2 promoter and the coding region was amplified from the genomic DNA of cells in line 1 after 50 transfers.

Complementation of ACE2 was performed by cloning the promoter and the coding region from each of the six lines after 60 transfers on the plasmid pT1-3B and transforming the resulting plasmid in an ACE2Δ mutant. The ACE2 plasmids from each of the six lines are called pACE2L1, pACE2L2, pACE2L3, pACE2L4, pACE2L5, and pACE2L6. The promoter and the coding region from each of the six lines was amplified using the primers given above. The region was amplified from DNA isolated from a single colony on YPD plates from each of the six lines.

### ACE2 knockout

The strain ΔACE2 was constructed by replacement of the ACE2 coding region with a hygromycin B resistance gene. For this, the gene HPH was amplified using the template pUG75, with primers 5’ATG GAT AAC GTT GTA GAT CCG TGG TAT ATA AAT CCC TCA GGC TTC GCG AAG CAG GTC GAC AAC CCT TAA T3’ and 5’TCA GAG AGC ATC AGT TTC GTT TGA AAG GGT GCG GTT CGA GTT TTG CTC GTA GTG GAT CTG ATA TCA CCT A3’. The PCR product was transformed in the respective strain and single colonies selected for on plates containing hygromycin B at a concentration 200 μg/ml. The replacement of ACE2 with HPH was confirmed using primers 5’CTC AAG CAA CAG TTA AAG TGC3’ and 5’TGT TAC TAT TAT TTA TTA TG 3’.

### Sequencing of ACE2 gene

Genomic DNA was isolated from ancestor and six replicate populations from both forward and reverse evolution as described previously (43). ACE2 gene was amplified using forward (5’ATG GAT AAC GTT GTA GAT CCG TGG TAT3’) and reverse (5’TCA GAG AGC ATC AGT TTC GTT TGA AAG3’) primers and sequenced. The coding sequence of the gene AMN1 was amplified using the primers (5’ATT TAT CAT TTT CCT TTT CT3’) and (5’ TTA CTG GTG GTC TGT TGT GTA 3’). All sequencing was done by Eurofins Scientific.

## Results and discussion

### Evolution of multicellularity on serial propagation under centrifugal selection

Unicellular to multicellular transition of yeast was studied using settling as selection (24). Figure 2A shows the representative images of yeast multicellular cluster populations during centrifugal selection. By 60 transfers, we note complex multicellular clusters in replicate populations (although cluster sizes vary across different lines). The dynamics of the change in phenomenology between the different independent lines is largely identical with each other in four of the six lines; but not the other two. Such parallelism in evolutionary experiments has been reported in the past (44–46). It has been suggested that even under controlled environmental growth conditions, the evolution of multicellular clusters generates spatial structure with increasing localized interactions and creates new spatial gradients (19). The evolution of multicellular clusters is driven by both chance and environmental changes (24, 27).

**Figure 2.**
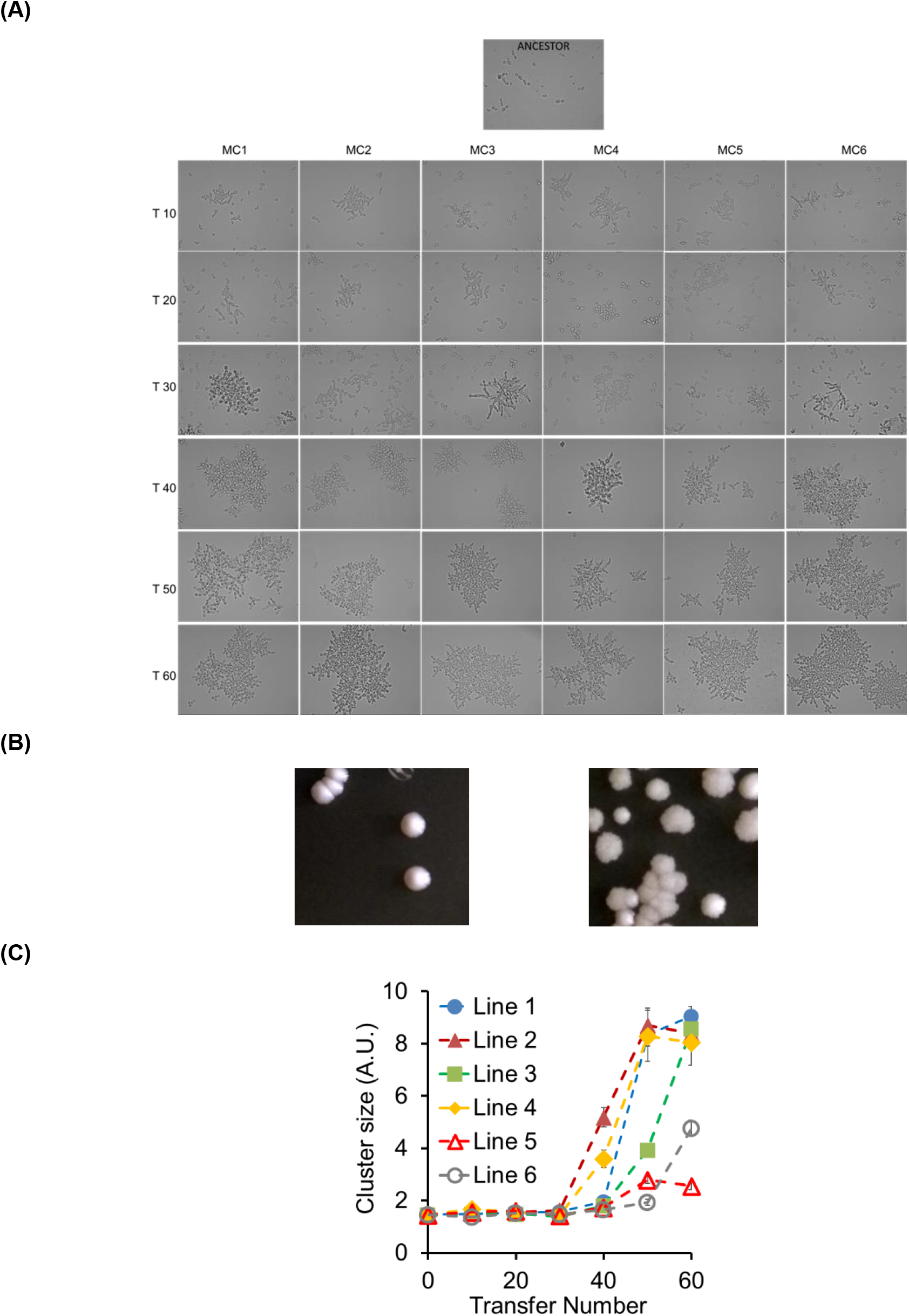
**(A)** The representative images of yeast multicellular cluster populations during centrifugal selection. An aliquot of overnight grown yeast cultures (both ancestor and multicellular cluster populations (MC1-MC6) from different transfer times (T10-T60) were placed on a microscopic slide covered with a cover slip. Images were taken using a microscope under 40X magnification. **(B)** Smooth (left) and rugose (rough) (right) morphologies of yeast unicellular and multicellular colonies, respectively. Overnight grown yeast cultures (both unicellular ancestor and multicellular cluster populations) were streaked on to YPD agar plates. Plates were allowed to incubate at 30°C until single colonies appear and images of the plates were then taken. **(C)** Dynamics of changes in yeast cluster size after 60 rounds of centrifugal selection. Exponentially growing yeast multicellular cluster populations (Line 1-6) were washed with PBS and analyzed for FSC-A of each cells/cluster using flow cytometer. More than 30,000 clusters per population were analyzed and the average of the FSC-A at the indicated time of transfer was calculated. The experiment was performed three times and the average and the standard deviation is reported.

When streaked on YPD agar plates, the evolved lines exhibit an altered colony morphology. These plates were incubated at 30°C until we see single colonies. As shown in Figure 2B, yeast ancestor and multicellular cluster colonies exhibited a distinct surface morphology. Yeast multicellular colonies showed rugose (rough) colony morphology, whereas unicellular ancestor showed a smooth morphology. This is a result of the cluster structure dictating the colony surface structure in the evolved lines.

### Dynamics of changes in yeast cluster size with serial propagation under centrifugal selection

Centrifugal selection favors the evolution of larger cluster size (24) and divergence in cluster size evolves rapidly during the transition from unicellularity to multicellularity (27). Therefore, in this study the average cluster size of yeast replicate populations was quantified as mean value of forward scatter area following every ten transfers using a flow cytometer. The size of the yeast clusters increased with concomitant increase in transfer time in most of the replicate populations (MC1-4 and MC6, but not MC5) (Figure 2C). While multicellularity evolved in all six replicate populations, it is also evident that multicellular clusters in the different lines did not evolve to the same size, and dynamics of evolution of the mean cluster size was not identical in multiple lines. Microbial evolution experiments have demonstrated that diversity may depend largely on environmental conditions (47) and ecological interactions between different genotypes (48).

Expectedly, the multicellular yeast clusters settles faster with increasing cluster size. It is evident from the Figure 3 that, four of the multicellular replicate populations (L1-L4) started settling faster after 40 transfers. However, replicate populations L5 showed moderate settling even after 60 transfers. The first snowflake yeast clusters formed are small during early transfer times, therefore they settle slowly. However, settling rate increased rapidly with the evolution of large cluster size during later stages of transfer times.

**Figure 3.**
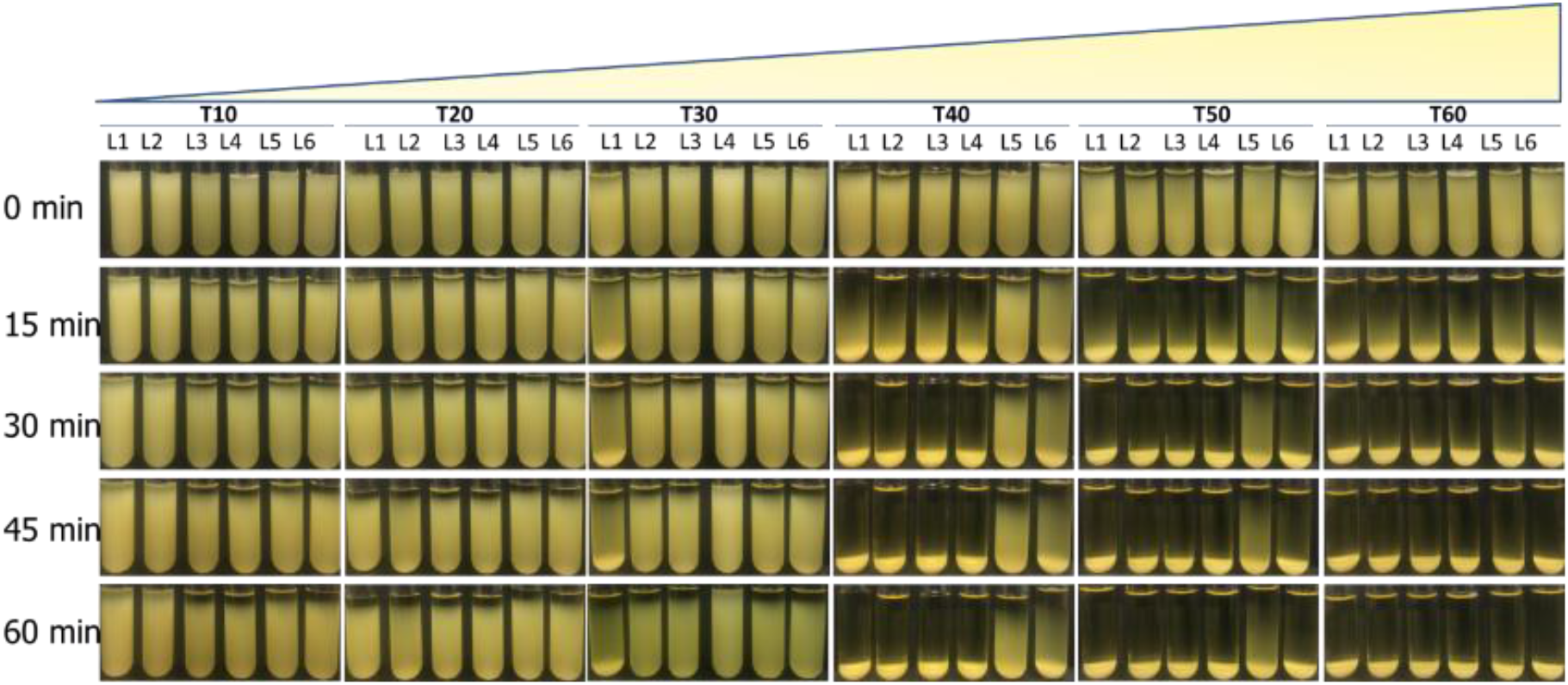
Yeast multicellular clusters settle faster with increase in transfer time. Six independent yeast lines (L1-L6) at different time points during forward evolution were inoculated from colonies into 10 mL YPD liquid cultures and grown overnight at 30°C. Each culture tube was vigorously vortexed and placed on a lab bench at room temperature to allow for settling. Images of each population during the settling were taken at different time intervals (0, 15, 30, and 60 min).

**Figure 3.**
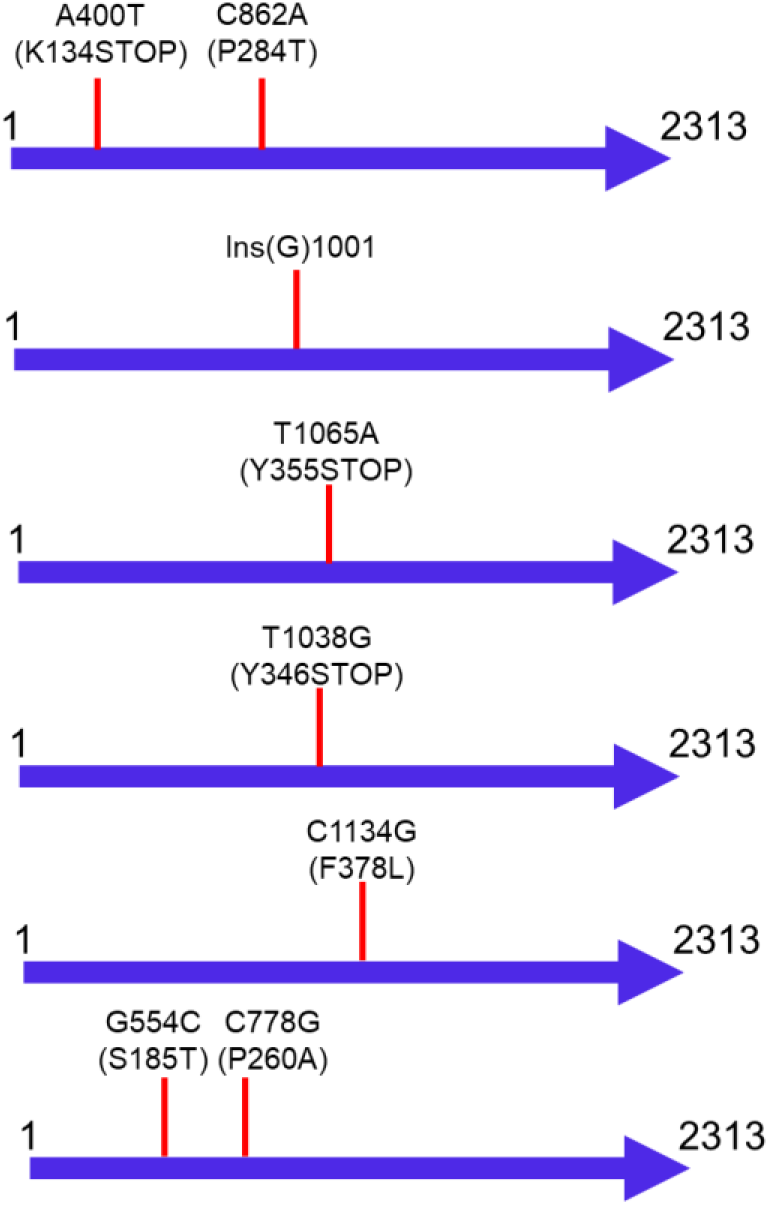
Mutations in ACE2 coding region in the six evolved lines after 60 transfers. The topmost figure is for Line 1, and the bottommost for Line 6.

### Multicellular clusters exhibit a tradeoff between growth rate and resistance to stress

Multicellularity is associated with multiple benefits such as reduced predation and division of labor between cells. However, these benefits come with costs. Multicellular clusters showed lower growth rates relative to their unicellular counterparts because of limitations of space and resource acquisition (27). The geometric constraints such as low surface area to volume ratio and of spatial organization in multicellular clusters limit the extent of interactions with the nutrient resources in the environment, and subsequently affects the metabolism, growth rate and viability (49). It has been suggested that there may be a trade-off between individual cluster size and growth rate (27). Therefore, we investigate whether cluster phenotype has any effect on the growth kinetics. As shown in Figure 4A and 4B, multicellular cluster populations grow slower when compared to their unicellular counterparts. As per the size-complexity rule, bigger entities have a greater division of labor than smaller ones (50).

**Figure 4.**
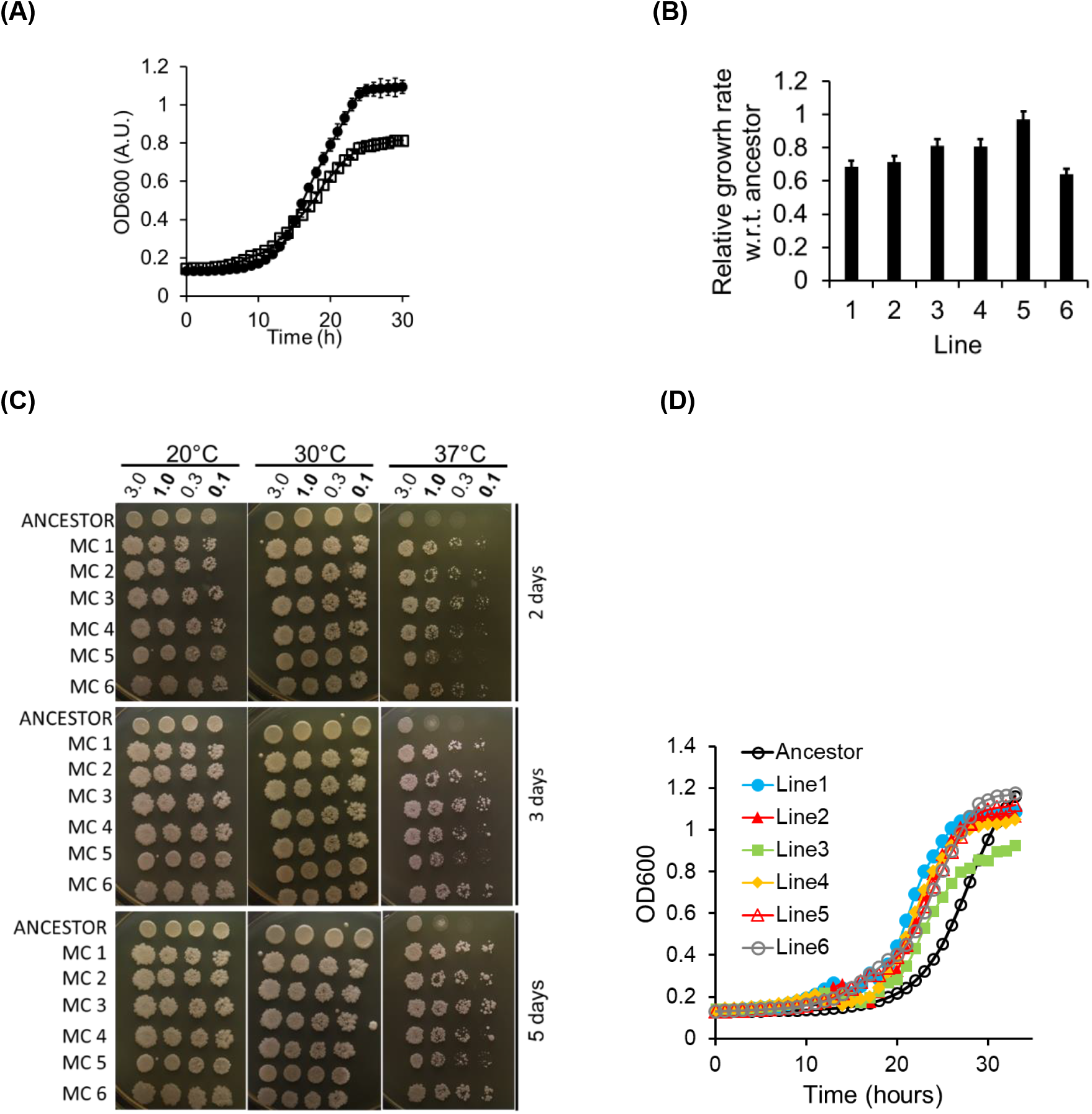
**(A)** Yeast multicellular cluster populations (empty square) show slow growth compared to unicellular ancestor (closed circles). Overnight yeast clusters and ancestor lines were diluted to an initial OD_600_ of 0.1 and the optical density was measured for every 1 h. The data represents average of six independent repeats. **(B)** Relative growth rate of yeast clusters with respect to ancestor. The experiment was performed in triplicate and the mean and the standard deviation are represented. **(C)** Yeast multicellular cluster phenotype confers heat stress resistance. Overnight yeast clusters and ancestor lines were diluted to OD_600_ of 3, 1, 0.3 and 0.1 and spotted on to YPD plates. Plates were incubated at indicated temperatures and images were taken at different days. MC=multicellular cluster population. **(D)** Effect of ethanol on the growth kinetics. Yeast multicellular cluster populations exhibit a shorter lag phase indicating resistance to ethanol stress compared to unicellular ancestor phenotype. Overnight yeast clusters and ancestor lines were diluted to an initial OD_600_ of 0.1 and exposed to ethanol (2.5 percent). Then the optical density was measured for every 1 h. Data is shown from the average of the six independent lines. Note that the shorter lag of the cluster’s growth, when compared to the ancestor, is not present when the cultures are grown in YPD (Figure 2A).

**Figure 4.**
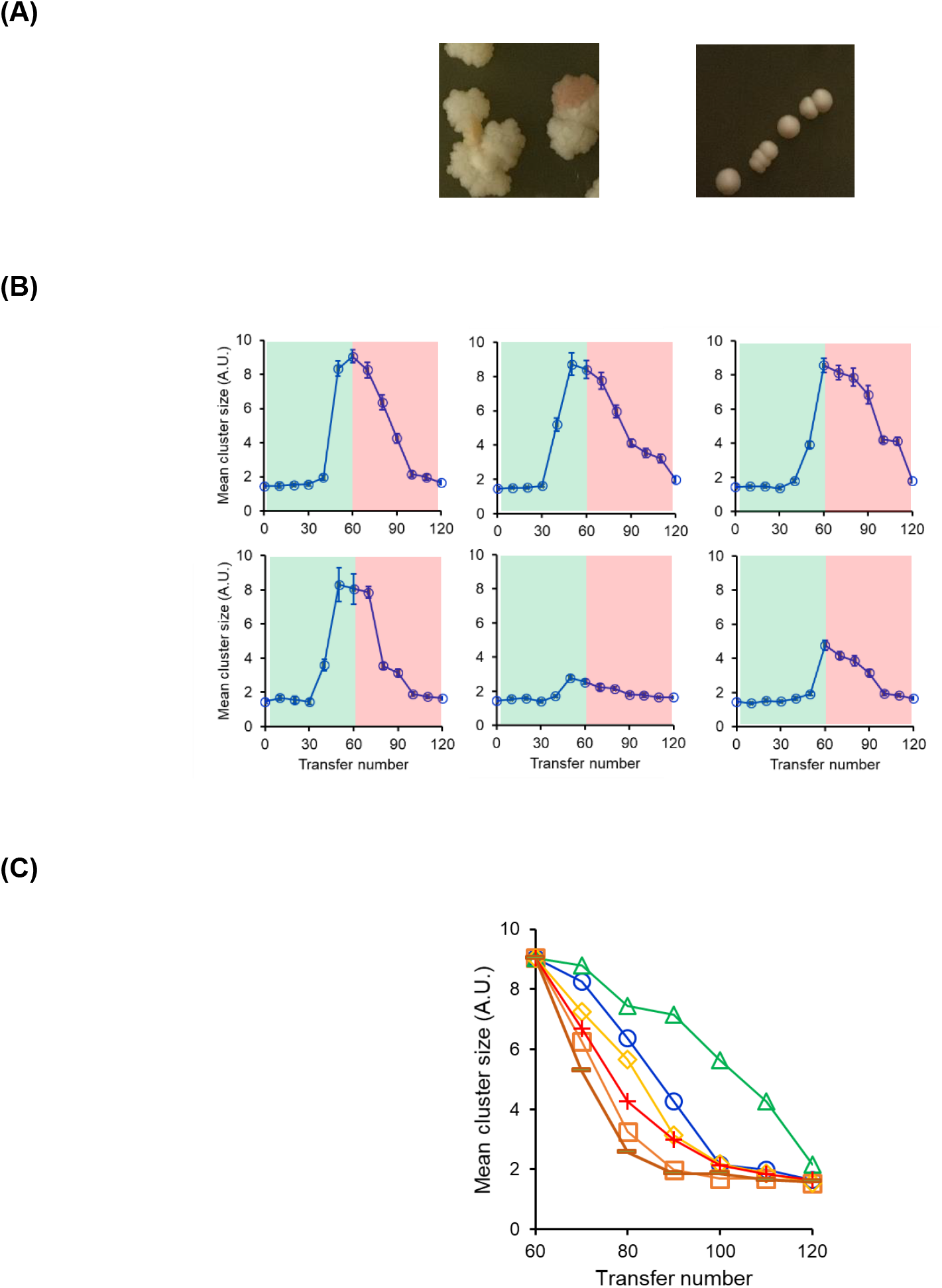
**(A)** Reversion of rough colony morphology to smooth colony morphology during reverse evolution from multicellularity to unicellularity. Overnight grown yeast cultures (both multicellular cluster population and population evolved from it during reverse evolution) were streaked on to YPD agar plates. Plates were allowed to incubate at 30°C until single colonies appear and images of the plates were taken. Images from the multicellular line 1 (left) and reversed evolved Line1 (right) are shown. **(B)** Dynamic trajectory of evolution of multicellularity (0-60 transfers) and evolution of unicellularity (60-120 transfers). Green shows transfers 0-60 when lines are evolved for multicellularity. Red shows transfers 60-120, when lines are evolved for unicellularity. The top row shows data for Lines 1-3 (left to right) and the bottom row for Lines 4-6 (left to right). The trajectories of evolution of multicellularity and that of its reversal are statistically distinct from each other for lines 1, 2, 3, and 6 (P<0.1, unpaired t-test). **(C)** Cluster size for 6 independent repeats of reverse evolution of multicellular clusters from Line 1. Five out of the six trajectories are statistically different from the trajectory of evolution of multicellularity (P<0.1, unpaired t-test).

A transition to cluster form confers selective advantage over unicellularity in ecological niches (17, 51). Examples of this phenomenon include protection from phagocytosis (52), predators (53), environmental stressors (ethanol and antibiotics) (54) and uptake of harmful compounds (55). Clustering may be beneficial for more efficient exchange of external resources between component cells of a cluster by reducing interactions with non-cooperative individuals (or cheaters) (i.e. increased efficiency of cooperative feeding) (26). It has also been suggested that cluster levels selection is beneficial for the use of growth-promoting secretions as well as exclusion of cheaters (19). This division of labor can allow multicellular organisms to perform tasks more efficiently, and can facilitate the evolution of complex traits (1). We study the heat and solvent stress resistance of yeast clusters and compare it with the unicellular ancestor’s performance in these environments.

#### Heat stress

Previous studies have reported that wild type yeast clusters strains were resistant to free/thaw treatment (31). We use spot assay to investigate the sensitivity of yeast cluster phenotype to cold (20°C) and heat stress (37°C). Multicellular yeast cluster lines (MC 1-6) were found to be resistant to heat stress (37°C) when compared to unicellular ancestor (Figure 4C). Even after five days of incubation, there is no significant growth of ancestor at indicated cell densities viz at OD600 of 0.3 and 0.1. Cold stress has no effect on the survival of both unicellular and multicellular cell lines.

#### Solvent stress

Toxic effect of ethanol on yeast is mainly caused by damaging the mitochondrial DNA and inactivation of enzymes such as hexokinase and dehydrogenase (56). Yeast multicellular clusters showed longer lag phase when compared to unicellular ancestor (Figure 4D). None of the cultures grew on higher ethanol concentrations (5% and 10%) (Figure 5).

**Figure 5.**
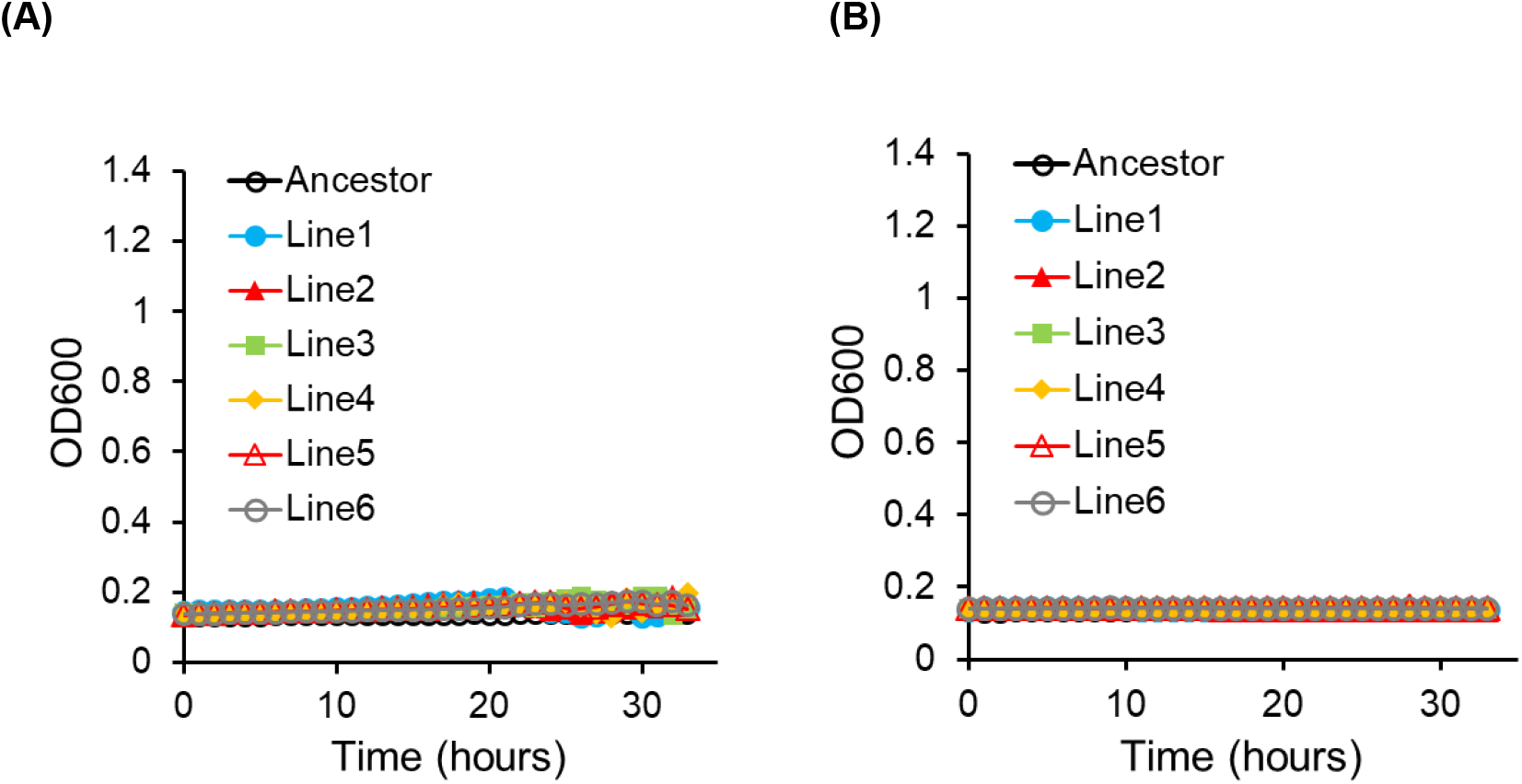
Growth curves of the ancestral and evolved lines in (A) 5% ethanol, and (B) 10% ethanol.

### Mutations in ACE2 transcription factor leads to evolution of the multicellular phenotype

Transition to multicellularity is typically associated with increases in the numbers of genes involved in three important processes such as cell differentiation (57), cell-cell communication (58, 59), and adhesion (17, 60, 61). The genetic basis for the evolution of multicellular cluster phenotype in yeast is due to disruption of ACE2 gene function. It has been reported that early (7-day) snowflake yeast clusters showed down-regulation of seven genes (CTS1, DSE4, DSE2, SUN4, DSE1, SCW11 and AMN1) which are important for daughter cell separation. Many of these genes act directly to degrade the bud neck septum. Previous studies have shown that the mutation or knockout of ACE2 gene prevents daughter–mother cell separation results in the formation of the snowflake phenotype (33, 62, 63). These experiments demonstrate that a single genetic mutation not only can create a new level of biological organization, but can also facilitate the evolution of higher-level complexity (34). Ratcliff et al reported non-synonymus mutations in that five of the ten replicate populations studied. In our experiments, all six lines acquired mutations in the ACE2 gene.

All six lines in our experiment, after 60 transfers, had mutations in the coding region of the gene (Figure 6). Line 1 had a SNP at position 400, which changed Lysine residue to a STOP codon and a non-synonymous change at position 862 (C to A; Pro to Thr). Line 2 had an insertion of nucleotide ‘G’ at position 1001 in the coding region of ACE2. Line 3 had a SNP (T to A) at position 1065, which changed a Tyrosine residue to a STOP codon. Line 4 had a T to G SNP at position 1038 in the coding region. This SNP changed a Tyrosine to a STOP codon. Thus, no functional copies of ACE2 were being produced in the cell in these four lines. Line 5 had a SNP at position 1134 in the coding region of ACE2 (C to G). This lead to a non-synonymous change in the coding region of the protein (Phe to Leu). Line 6 had SNPs at positions 554 (G to C; Ser to Thr) and 778 (C to G; Pro to Ala). To verify the effect of the mutant allele of ACE2 in dictating the multicellular cluster, the ACE2 promoter and the coding region from each of the six lines were transferred to an ACE2Δ mutant. Transferring the ACE2 allele lead to the multicellular phenotype in each of the six complementation experiment (Figure 7). We do not discount the possibility of other molecular determinants dictating the evolution of the multicellular cluster. However, our results show, mutations in ACE2 is a key molecular determinant in dictating the evolution of the multicellular phenotype. Sequencing of the ACE2 region in line 1 after 40 and 50 transfers revealed that the non-synonymous mutation (P to T) was acquired first. This mutation was first observed after 40 transfers. Thereafter, the A400T mutation was observed after 50 transfers. Thus, the increase in cluster size was concomitant with the accumulation of mutations at the ACE2 locus. The coding region of AMN1 was also sequenced in all six lines after 60 transfers. However, no mutation was identified in this region of the chromosome.

**Figure 6.**
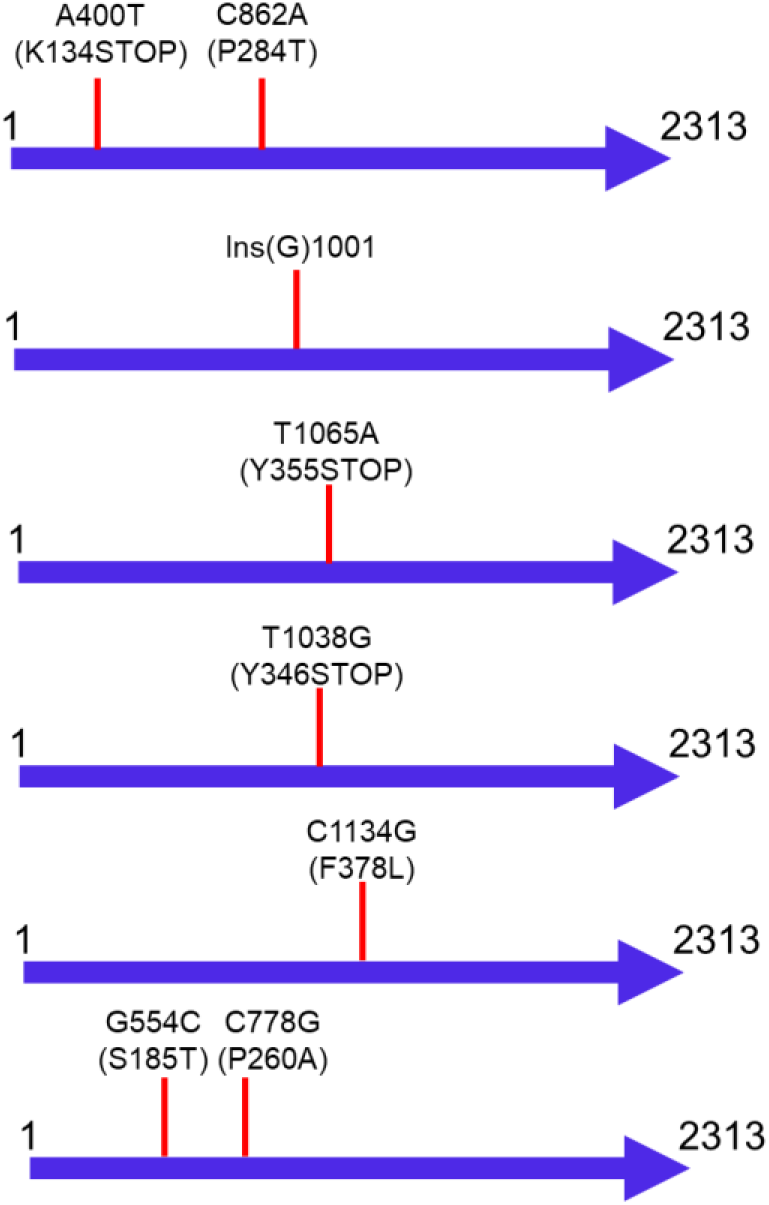
Mutations in ACE2 coding region in the six evolved lines after 60 transfers. The topmost figure is for Line 1, and the bottommost for Line 6.

**Figure 7.**
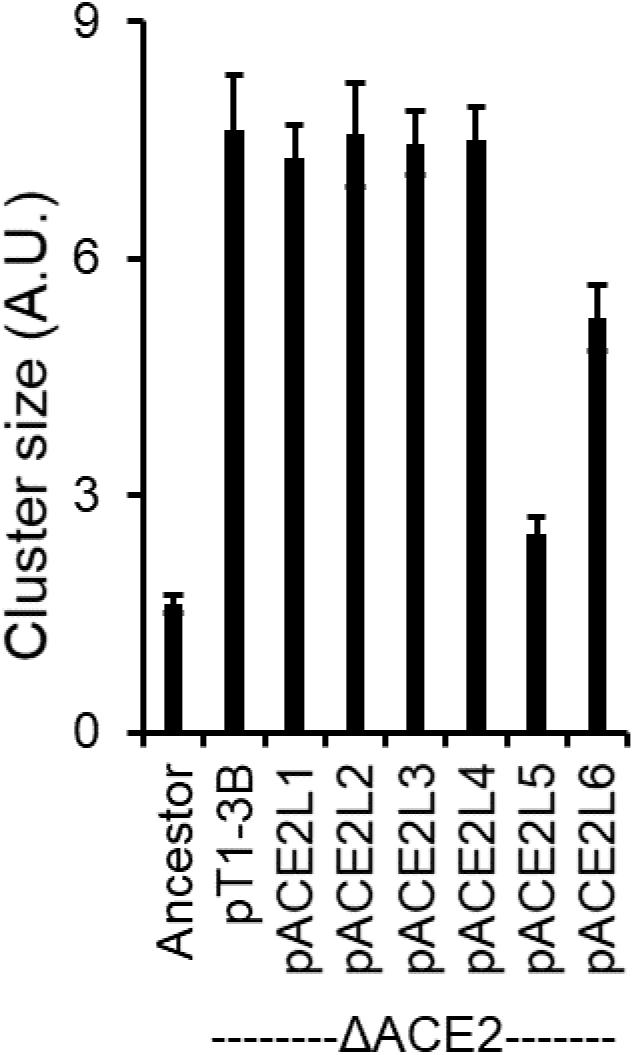
ACE2 mutant alleles from the evolved lines, complementing in an ACE2Δ deletion haploid strain.

### Reversal of centrifugal selection leads to evolution of unicellularity from multicellular clusters

Compared to unicellular ancestor, multicellular clusters grow slower in culture media. The growth limitation is due to increased competition and mass transfer limitations regarding access to resources (20). Further, their complexity makes them more vulnerable to disruption (1). Given this, in what contexts can we study evolution of unicellularity, starting with a multicellular cluster?

In the present study, we aimed at investigating the reverse evolution from multicellularity to unicellularity using experimental evolution, under the same centrifugal selection pressure. Growing the multicellular cells the same way, and after the same processing, the cells from the topmost layer of the liquid were transferred to the next tube. In this way, the selection pressure in the experiment was exactly the opposite of the one applied in the uni- to multicellular evolution. This reverse evolution experiment leads to several physiological changes. Firstly, the average cell size in the clusters decreased rapidly and unicellular lines were evolved. As a result, the colony morphology was restored from rugose to smooth colony morphology during reverse evolution from multicellularity to unicellularity (Figure 8A). It has been suggested that both genotypic and phenotypic factors can affect the colony morphology. Some cellular properties, such as cell adhesion, and budding pattern, have also been associated with morphological changes in *S. cerevisiae* (64).

**Figure 8.**
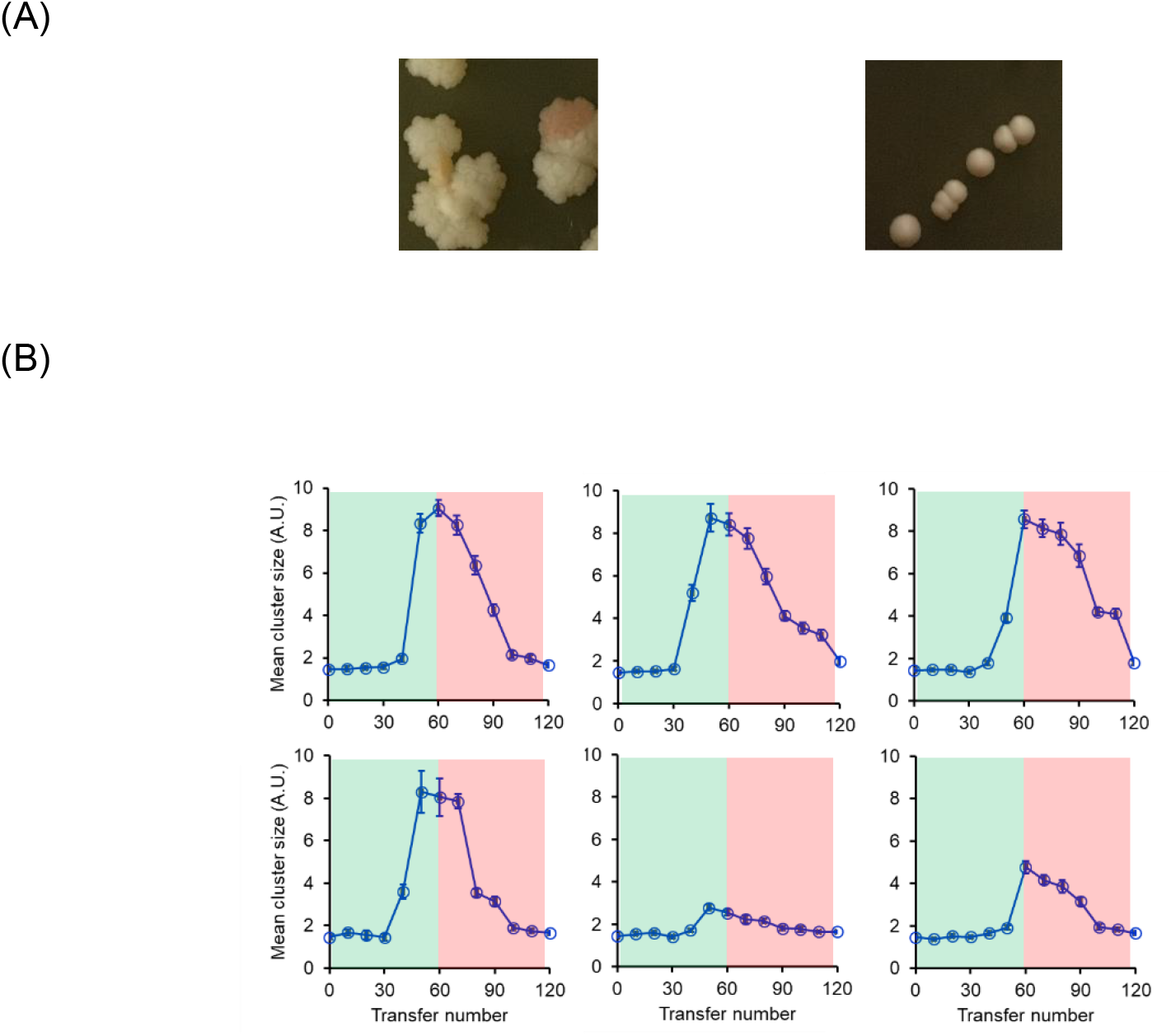
(A) Reversion of rough colony morphology to smooth colony morphology during reverse evolution from multicellularity to unicellularity (one of the six lines is shown). Overnight grown yeast cultures (both multicellular cluster population and population evolved from it during reverse evolution) were streaked on to YPD agar plates. Plates were allowed to incubate at 30°C until single colonies appear and images of the plates were taken. **(B)** Dynamics of trajectories during evolution of multicellularity (green) and during evolution of unicellularity (red). Top row (left to right) is data for lines 1-3. Bottom row (left to right) is data for lines 4-6.

The trajectory of the cluster size in the reverse evolution is shown in Figure 8B. As shown in the Figure, in most of the lines, the transition from multicellular to unicellular phenotype (transfers 60 to 120) has a trajectory distinct from the original transition from unicellular to multicellular phenotype (transfer 0 to 60). This data suggests that the evolutionary trajectory does not retrace its path, at a phenotypic level. Second, the growth kinetics of the reverse evolved lines was restored closer to that of the ancestral strain (Figure 9). This is likely due to the transport limitations being eliminated in the reverse evolved unicellular lines.

**Figure 9.**
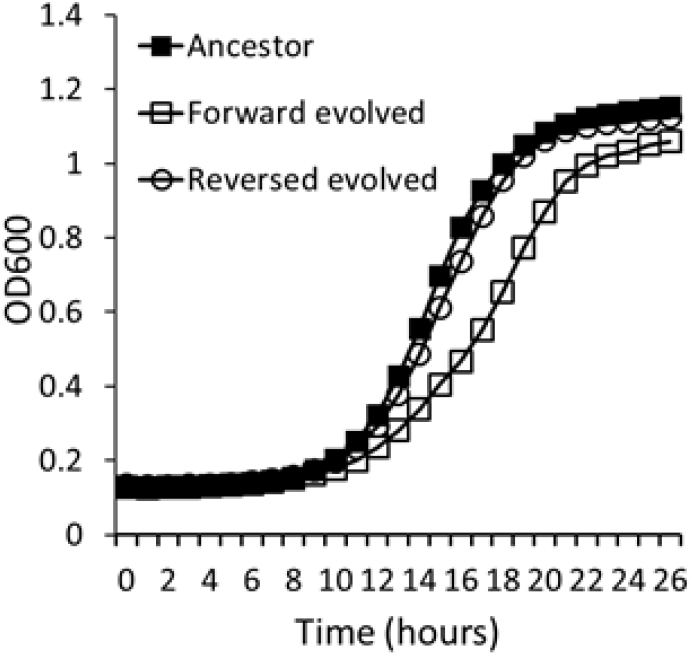
Growth kinetics of ancestor, forward-, and reverse-evolved line. Data for one of the six lines is shown. The data is average for six independent experiments. The standard deviation for the data is less than 3% of all data points.

To understand the molecular basis which trigger the reverse evolution, after 60 transfers, we sequenced the ACE2 region of the reverse evolved lines. However, none of the four lines had acquired a mutation in the ACE2 region, when compared to the corresponding sequence of the multicellular cluster that each line had started from. Thus, the evolutionary transition from the multicellular to the unicellular phenotype is brought about by mutations elsewhere on the chromosome.

Additionally, six independent lines from the multicellular clusters from line 1 were reverse evolved, selecting for unicellularity. The premise behind this experiment was to test for repeatability of evolution and identify paths where the phenotypic trajectory is re-traced. Our results demonstrate that significant differences exist between the evolutionary paths traced by the six independent lines in the reverse evolution (Figure 10). Moreover, none of the six paths is statistically identical with the path traced when the line was evolved for multicellularity. Sequencing of the ACE2 coding region revealed that none of the six independent lines in the reverse evolution had acquired a mutation. Thus, the molecular determinants of the reverse evolution experiment lie elsewhere on the chromosome.

**Figure 10.**
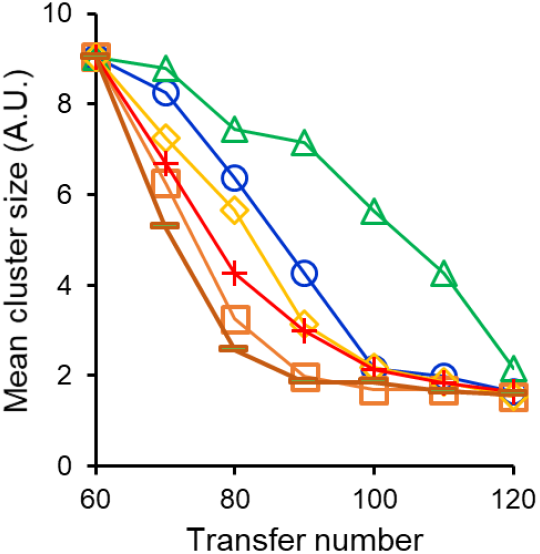
Trajectory of evolution of unicellularity starting from the multicellular cluster population in Line 1. Six independent repeats of reversal of multicellularity evolution exhibit distinct kinetics of reduction in cluster size. Five of the six trajectories are distinct from reversal of the original unicellular to multicellular cluster evolution (P<0.1, unpaired t-test).

## Conclusion

In this work, we report forward and reverse evolution between unicellularity and multicellularity in the yeast *S. cerevisiae*. In this context, selection of settling under gravity or centrifugation provides an environment where study of evolution, when selection is reversed can be studied. As has been reported previously, multicellularity readily evolves under the appropriate selection pressure. Of our six lines, four of them exhibit dynamics of cluster size increase quite close to each other. To that effect, the evolution of multicellularity is a repeatable and predictable process. Somewhat more surprisingly, all six lines acquire mutation at a single locus in the genome. This single set of mutations in the ACE2 gene is responsible for exhibition of the multicellularity trait (Figure 6). The mutations at this locus suggest that the selection coefficient associated with an ACE2 mutation is very high.

On reversal of the selection pressure, all six lines exhibit reversal of the multicellular phenotype. While the phenotype is completely reversed in these lines, the trajectory of the change in the cluster size does not mimic the forward evolution experiment. Moreover, sequencing the ACE2 region in the reverse evolved lines tells us that the unicellular phenotype did not evolve because of additional/reverse mutations in ACE2. Thus, the molecular basis of the reverse evolution experiment is at another locus/loci on the genome.

Our results also show that although the selection pressure of gravity leads to evolution of multicellularity, the multicellular phenotype confers advantages in the face of several environmental stresses too. This advantage has been observed in a number of studies (26, 65–67). Thus, in the context of evolution of multicellularity during the evolution, any of these stresses could provide an avenue for evolution of multicellularity, which is known to have evolved a number of times in the history of evolution of life on Earth (15). Reverse evolution from multicellularity to unicellularity has been studied in two different contexts (35, 68). In the first, natural isolates which formed multicellular clusters were reverse evolved for unicellularity. The authors report that mutations at the AMN1 (69) locus were responsible for evolution of unicellularity. The dynamics of this transition was, however, not reported. In a second study, reversal to unicellularity was evolved via propagation on solid media, in a structured environment. The selection forces driving this transition are, hence, not the opposite of those which drove evolution of multicellularity.

From our reverse evolution experiment, we note that the dynamic trajectory of reverse evolution is different from that of the forward one. In the context of reverse evolution, what is the genetics at the organismal level, which constrains this evolutionary process? (70, 71). If we assume that the strength of selection is reversed exactly in the reverse experiment, then what could lead to the difference in the trajectories? Several factors could be responsible for this. First, with the evolution of multicellularity, the size of the population in the culture media changes. Population size is an important determinant of the rate of adaptation (72). Second, mutation rates across the genome are known to be variable (73). Hence, since in this case, the molecular determinants of the forward and reverse evolution are different, these might be subject to different mutation rates. Third, the number of adaptive mutations facilitating forward and reverse evolution might be different, and as a result, the time of transition from one state to the other may be different. This is especially so since the genetic backgrounds in which forward and reverse evolution of multicellularity is being studied are different from each other. In this context, what role, if any, does epistasis play in this process is also not known and can be a determinant in dictating the evolutionary trajectory of the cluster size (74). Finally, how cluster size relates with the accumulation of mutations is not yet known. The relationship between these two variables in the forward and reverse evolution experiment need not be the same. Determining the answers to these questions related to the forward and backward adaptive processes forms the basis of the future work in this direction.

## Acknowledgements

PA was funded by the IITB Institute Post-Doctoral Fellowship. AM is supported by the Council of Scientific and Industrial Research (CSIR), Government of India, as a Senior Research Fellow (09/087(0873)/2017-EMR-I).

